# Enhancing Adult Neurogenesis Rescues Hippocampal Cognitive Functions in an Alzheimer’s Mouse Model

**DOI:** 10.64898/2026.03.25.713986

**Authors:** Chi-Chieh Lee, Federico Calegari

**Affiliations:** Center for Regenerative Therapies Dresden (CRTD), Technische Universität Dresden, Dresden, Germany; Mortimer B. Zuckerman Mind Brain Behavior Institute, Columbia University, New York, United States

## Abstract

Alzheimer’s disease (AD) is the most prevalent form of dementia, characterized by progressive memory loss, cognitive decline, and emotional dysregulation. Adult hippocampal neurogenesis (AHN) critically contributes to cognition and mood but undergoes precipitous decline during AD progression. Here, we investigated whether enhancing AHN through genetic expansion of endogenous neural stem cells (NSC) ameliorates AD-related phenotypes. Using lentiviral overexpression of the cell cycle regulators Cdk4 and CyclinD1 in the dentate gyrus of 3xTg-AD mouse, we show that enhancing AHN partially rescues hippocampal-specific cognitive functions, namely: spatial navigation and exploratory behavior. These findings show that endogenous NSC can be exploited to ameliorate hippocampal cognitive functions in AD, providing additional evidence for exploiting AHN as a promising therapeutic target for neurodegenerative disease.

## INTRODUCTION

Alzheimer’s disease (AD) is a progressive neurodegenerative disorder characterized by profound memory loss and cognitive decline. The pathological cascade underlying AD involves the accumulation of amyloid-beta (Aβ) plaques and tau tangles, leading to neuronal loss and disruption of neural networks that result in memory impairments and mood dysregulations (Breijyeh and Karaman, 2020; Knopman *et al*., 2021). The hippocampus, a critical hub for memory processing, spatial navigation, and emotional regulation (Burgess, Maguire and O’Keefe, 2002; Strange *et al*., 2014; Eichenbaum, 2017; Moser, Moser and McNaughton, 2017), represents a primary target of AD pathology and focal point for therapeutic intervention.

As a hallmark of the hippocampus, an endogenous reserve of neural stem cells (NSC) resides within the dentate gyrus (DG), serving as a source of adult-born neurons throughout life in a process referred to as adult hippocampal neurogenesis (AHN). AHN plays a pivotal role in maintaining the plasticity and adaptability of the hippocampal circuitry (Toda *et al*., 2019). Highlighting its multifaceted role, AHN not only positively contributes to learning and memory processes but is also linked to mood regulation (Anacker and Hen, 2017; Kempermann, 2022). These roles of AHN are particularly relevant, providing avenues to rescue deficits resulting from AD with significant clinical implications. However, as one of the earliest features of AD, AHN undergoes a precipitous decline during disease progression, raising concerns pertaining to its significance and therapeutic potential (Moreno-Jiménez *et al*., 2019; Salta *et al*., 2023).

Given AHN’s critical roles in cognition and mood, multiple strategies have emerged to enhance neurogenesis as a therapeutic approach for AD. These strategies include stimulation of endogenous AHN through physical exercise or other physiological stimuli (Choi *et al*., 2018; Valenzuela *et al*., 2020). While partially effective, these approaches could not discriminate between effects triggered by AHN within the DG versus any of the many systemic effects that the physiological stimuli themselves are known to trigger, including massive re-wiring of the whole hippocampal connectivity and activity (Emery *et al*., 2023). Alternative strategies to attempt to rescue neuronal loss included cell therapy interventions involving NSC transplantation (Si and Wang, 2021) or glia-neuron reprogramming (Zhou *et al*., 2022), which in either case was ineffective in the replacement of neurons undergoing physiological maturation, specification and functional integration. Recent molecular interventions utilizing microRNA or preventing neuronal death have provided stronger evidence for a role of AHN in the rescue of AD (Walgrave *et al*., 2021; Mishra *et al*., 2022). Nonetheless, additional proof-of-principles are important to validate the effects of enhancing endogenous NSC, without systemic effects or external cell sources, to restore hippocampal function in an AD-compromised neurogenic niche. This is critical for establishing AHN as a robust and reproducible therapeutic strategy for AD.

In this context, our group has previously developed a viral transduction approach based on the overexpression of the cell cycle regulators Cdk4 and CyclinD1 (together referred to as 4D) (Artegiani, Lindemann and Calegari, 2011) that alone is sufficient to promote the expansion of endogenous NSCs. This approach resulted in the generation and functional integration of 2-3 fold higher number of newborn neurons in the mouse DG over the course of life and without exhaustion of NSC (Berdugo-Vega *et al*., 2020), which was accompanied by improvements in cognitive flexibility and memory indexing (Berdugo-Vega *et al*., 2021). It is noteworthy to mention, however, that our previous results were obtained in a healthy neurogenic. Whether 4D could still be effective in promoting NSC expansion and compensate cognitive decline in the AD-compromised hippocampus is unknown. To this aim, here we investigated the efficacy of the 4D system in enhancing AHN within the 3xTg AD mouse model and assessed its impact on hippocampal cognitive function.

## RESULTS

### 4D overexpression rescues AHN in 3xTg-AD mice

Our group has previously developed an approach to genetically increase the expansion of endogenous NSC by overexpression of the cell cycle regulators Cdk4 and cyclinD1 (together: 4D) (Artegiani, Lindemann and Calegari, 2011). Promoting AHN, injection of 4D lentiviruses in the DG was found to improve cognitive function, navigational performance, memory precision and indexing over the course of life (Berdugo-Vega et al., 2020; Berdugo-Vega et al., 2021). However, since previous studies were performed in healthy mice, it is unknown whether the same 4D strategy could be used to increase AHN and improve hippocampal function in an AD-compromised neurogenic niche. More specifically, mouse models of AD are characterized by a major reduction in the number of NSC and impaired AHN (Mu and Gage, 2011; Salta *et al*., 2023). These deficits arise early in life, in juvenile mice, well before the onset of amyloid plaque formation and cognitive impairments (Liu *et al*., 2023). In turn, this indicates that AD underlies a highly dysfunctional neurogenic niche in which 4D may not be sufficient to overcome barriers to rescue deficits in NSC numbers, their proliferation and/or survival of newborn neurons.

To explore the efficacy of our 4D approach during the progression of AD, we used the 3xTg AD mouse model. This model is widely used and closely resembles AD pathologies (Oddo *et al*., 2003), including cognitive deficits (Billings *et al*., 2005). Aligned with studies characterizing 3xTg AD in female mice (Clinton *et al*., 2007; Sterniczuk *et al*., 2010; Javonillo *et al*., 2022), we also observed cognitive deficits in the MWM arising at 6 months of age including a reduction of hippocampal-specific, allocentric navigation during learning by and increased perseverance upon reversal (change in probability of WT vs. AD: contextual learning = 171%, perseverance = 313%; logistic regression, odds ratio (OR) of WT vs. AD: contextual learning = 5.22, perseverance = 5.6; Wald-test *p* < 0.0001 and *p* < 0.001, respectively; Fig. S1A, details below). Also aligned with studies using this AD mouse model, we observed major increases in amyloid plaque formation progressing until the last point analyzed at around 10 months (immunoreactive area in hippocampus in healthy control and 3xTg AD, mean ± SD: 0.05 ± 0.04% vs. 0.43 ± 0.22%, *p* < 0.01; Fig. S1B). Next, to investigate the possibility to increase the expansion of endogenous NSC, 4D stereotaxic viral injections were performed targeting the DG of 6-month-old 3xTg mice (hereafter referred to as AD/4D). In parallel, age-matched wildtype or 3xTg AD mice injected with GFP viruses were used as healthy or experimental controls, respectively (hereafter referred to as WT/GFP or AD/GFP; Fig. 1A; top). Neither AD/GFP nor AD/4D mice showed any change in amyloid plaque burden (mean ± SD: 0.43 ± 0.22% vs. 0.51 ± 0.2%, *p* = 0.41; Fig. S1B).

**Figure 1.**
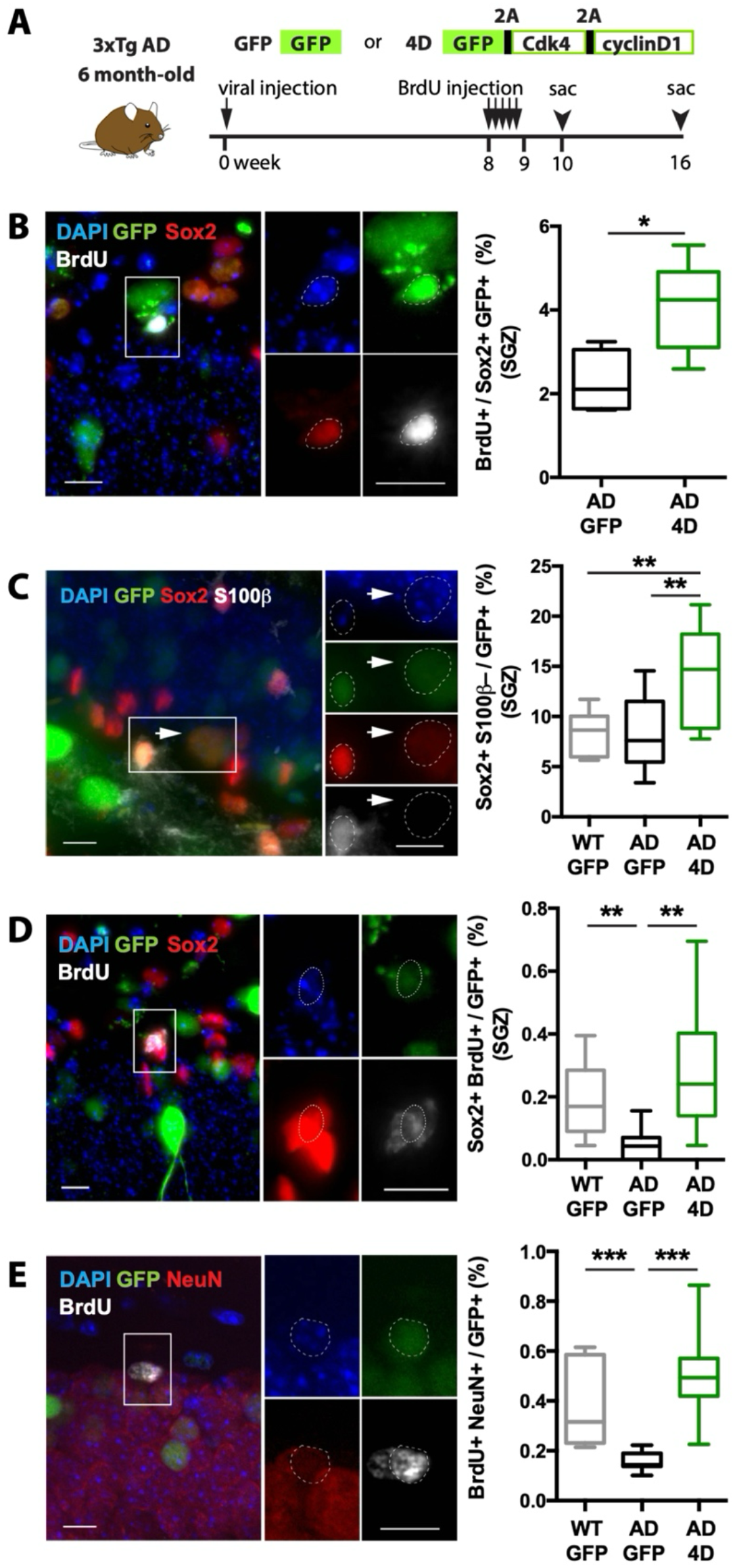
4D increases NSC expansion and neurogenesis in 3xTg AD mice. **A**) Scheme depicting the experimental layout to assess AHN upon injection of GFP or 4D viruses. **B**–**E**) Examples of fluorescence pictures (left) and quantifications (right) of cellular markers (as indicated) in subgranular zone (SGZ) or the DG of mice injected with GFP or 4D viruses (as indicated). Insets (dashed boxes) are magnified and examples of cells scored indicated (arrowheads or dotted lines) of brains harvested either 10 (B, n = 4 and 5 for AD/GFP, and AD/4D, respectively) or 16 (C-E, n = 11, 10, and 10 for WT, AD/GFP, and AD/4D, respectively) weeks upon viral injection. Values are represented as whisker box (see Table 1 for mean ± SD), unpaired Student’s t-test, * *p* < 0.05; ** *p* < 0.01, *** *p* < 0.001. Scale bar = 10 μm.

To validate the efficacy of 4D in the dysfunctional AD niche, we first examined the short-term proliferative responses of NSC by administrating 5-day BrdU followed by assessment of BrdU+Sox2+ infected cells 1 week thereafter. This showed a doubling in the proportion of proliferating NSC in AD/4D (AD/GFP vs. AD/4D: mean ± SD: 2.27 ± 0.76% vs. 3.97 ± 0.99%; *p* < 0.05; Fig. 1B). After validating the effect of 4D to promote the proliferation of NSC in the dysfunctional AD niche, we next attempted to recapitulate in a single experiment the cumulative long-term effects of diverse biological processes related to neurogenesis. To this aim, we performed BrdU labeling of proliferating NSC by a 5-day administration of BrdU 8 weeks after viral injection as above but this time sacrificing animals 8 weeks thereafter (Fig. 1A). This paradigm was chosen to infer, among the viral-transduced (GFP+) cells, parameters related to i) NSC abundance, ii) label retention within quiescent NSC, iii) overall neurogenic output and neuronal survival.

**Table 1.** Neurogenesis level.

| Neurogenesis level |  |  |  |  |
| --- | --- | --- | --- | --- |
| Group | BrdU+ / Sox2+ GFP+ (%) | Sox2+ S100b- / GFP+ (%) | Sox2+ BrdU+ / GFP+ (%) | NeuN+ BrdU+ / GFP+ (%) |
| WT+GFP | N/A | 8.610 ± 2.072 | 0.1845 ± 0.1102 | 0.3840 ± 0.1619 |
| AD+GFP | 2.268 ± 0.756 | 8.279 ± 3.652 | 0.0430 ± 0.04894 | 0.1582 ± 0.03696 |
| AD+4D | 3.968 ± 0.988 | 13.97 ± 4.980 | 0.2765 ± 0.1942 | 0.5071 ± 0.1698 |

With regard to the first parameter, abundance of bona fide NSC (Sox2+S100β−/GFP+), irrespective of their proliferative state (i.e.: BrdU incorporation), did not significantly change across control WT/GFP and AD/GFP mice (mean ± SD: 8.61 ± 2.07% vs. 8.28 ± 3.65%, respectively; *p* = 0.79). In contrast, about 2-fold increase was found in AD/4D (mean ± SD: 13.97 ± 4.98%) relative to both WT/GFP and AD/GFP mice (both *p* < 0.01; Fig. 1C). This doubling in the abundance of NSC upon 4D overexpression extends the previous effect on short-term proliferation (see above, Fig. 1B) and is fully consistent with previous reports in either young or aged mice (Berdugo-Vega *et al*., 2020; Berdugo-Vega *et al*., 2021).

Having confirmed the 4D-effects on enhanced NSC proliferation and numbers, we next investigated whether these translated into changes in the abundance of label-retaining NSC as a long-term reservoir for neurogenesis. We found that the number of BrdU+Sox2+, label-retaining cells in WT/GFP mice was substantially increased, by over 4-fold, relative to AD/GFP controls (mean ± SD: 0.18 ± 0.11% vs. 0.04 ± 0.05%; *p* < 0.01). Intriguingly, this deficit in quiescent NSC in AD/GFP mice was rescued in AD/4D mice (mean ± SD: 0.04 ± 0.05% vs. 0.27 ± 0.19%; *p* < 0.01; Fig. 1D) to values comparable to, or even greater than, WT/GFP mice.

Finally, third, total neurogenic output and/or neuronal survival was quantified as the proportion of birthdated, BrdU+ cells expressing the neuronal marker NeuN and still detectable at 8 weeks post BrdU birth dating. Consistent with previous reports on the negative effect of AD on adult neurogenesis (Mu and Gage, 2011; Salta *et al*., 2023), we found that WT/GFP mice displayed 2-fold greater levels of BrdU+/NeuN+ adult-born neurons relative to AD/GFP mice (mean ± SD: 0.38 ± 0.16% vs. 0.16 ± 0.04%; *p* < 0.0005). Notably, AD/4D mice showed an essentially complete rescue in the levels of neurogenesis with a 3-fold increase relative to AD/GFP mice (mean ± SD: 0.51 ± 0.17% vs. 0.16 ± 0.04%; *p* < 0.0001) and similar to, if not even greater than, WT/GFP healthy controls (*p* = 0.11; Fig. 1E).

Together, our data show that NSC preserve their identity and potential to respond to neurogenic stimuli even in the dysfunctional neurogenic niche of AD mice. In turn, our data underscore the ability of 4D to enhance and fully rescue AHN in 3xTg AD mice providing the means to exploit the endogenous pool of NSC to attempt to rescue hippocampal cognitive deficits.

### Enhanced neurogenesis partially restores hippocampal-related behaviors of 3xTg AD mice

While improvements in NSC expansion and neurogenic output were encouraging, the critical question remains whether enhancing AHN translates into cognitive benefits in AD mice. To address this, we induced 4D overexpression as previously described and 12 weeks later subjected 3xTg mice to the OFT and MWM. These tests were chosen to assess baseline exploratory behavior in the absence of cognitive challenge, and spatial learning and memory as a cognitive navigational task, respectively (Fig. 2A, 2D).

**Figure 2.**
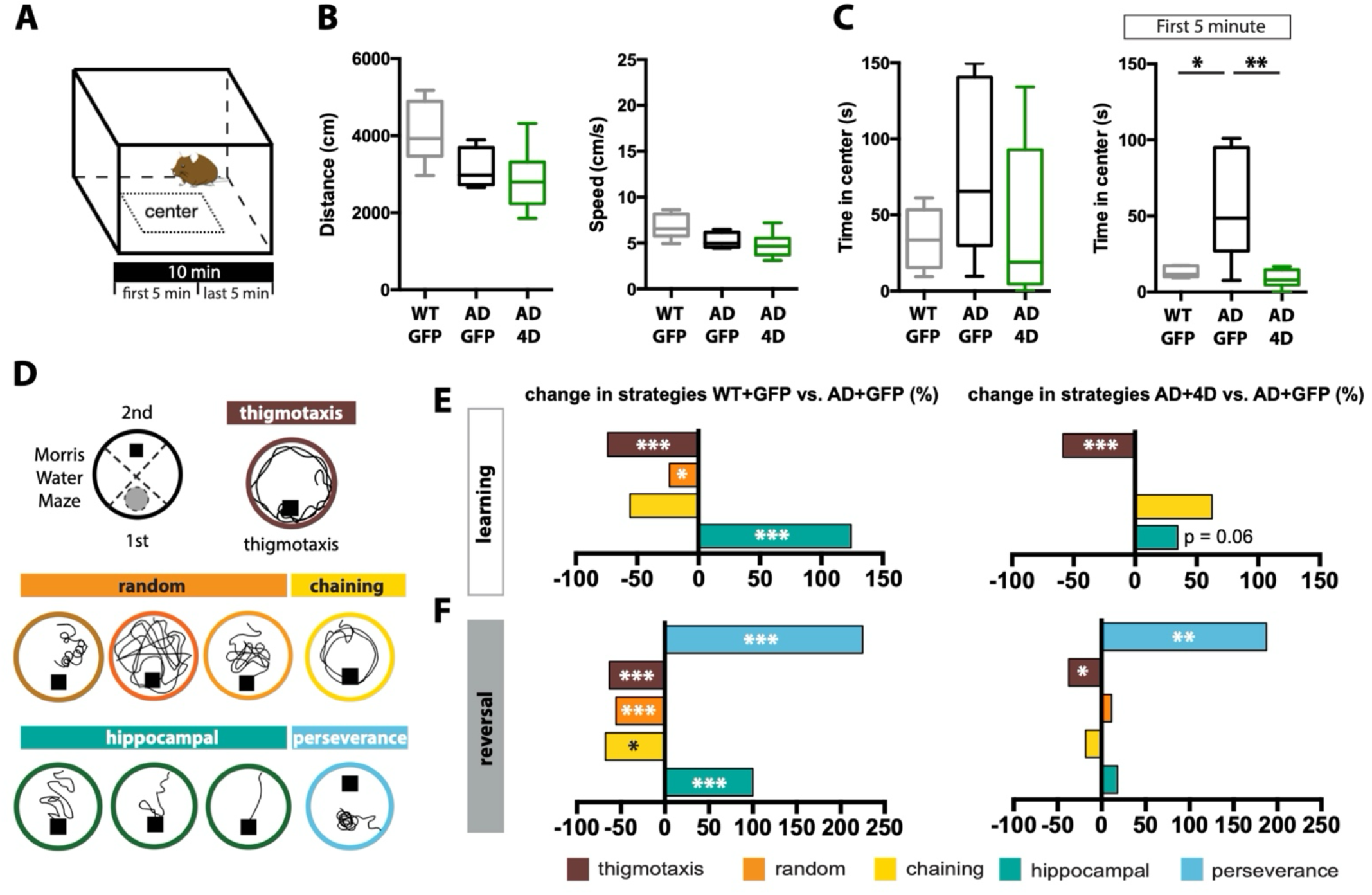
Enhancing neurogenesis rescues hippocampal deficits of 3xTg AD mice. **A-F)** Schematics of the OFT (A) and MWM navigational strategies (D). **B** and **C**) Distance (cm) and average speed (cm/s) during 10-minute exploration of the OFT (B) and time spent in the center (s) during 10- or 5-minute exploration (C). **E**) Comparison of the percentage change across trials during learning (top) or upon reversal (bottom) between WT and AD/GFP (left) or AD/4D and AD/GFP (right) mice. Values in (B) and (C) are represented as whisker box (n = 6, 5, and 6 for WT, AD/GFP, and AD/4D groups, respectively). Unpaired Student’s t-test, * *p* < 0.05; ** *p* < 0.01; *** *p* < 0.001. Values (see Table 2 for mean ± SD) in (E) and (F) represent strategy change (n = 11, 13, and 13 for WT, AD/GFP, and AD/4D groups respectively). Wald-test, * *p* < 0.05; ** *p* < 0.01; *** *p* < 0.001.

Overall, no difference in locomotor activity in the OFT was observed with regard to neither travel distance nor speed across groups of WT/GFP, AD/GFP or AD/4D mice (all differences *p* > 0.056; Fig. 2B). Independently from locomotor activity, we observed that during the entire test, 10 minutes in total, WT/GFP mice exhibited a trend of reduced exploration of the central area of the field relative to AD/GFP mice (mean ± SD: 34.3 ± 19.7s vs. 81.3 ± 58.1 s, *p* = 0.09; Fig. 2C, left). This increase in time spent within the central area displayed by AD/GFP mice was however not observed in AD/4D mice showing instead a not significant trend (AD/4D mean ± SD: 42.8 ± 52.8 s, *p* = 0.28; Fig. 2C, left) to reduce their exploration of the center to a degree more similar to healthy controls (AD/4D vs. WT/GFP, *p* = 0.72; Fig. 2C, left).

**Table 2.** Behavioral test results.

| Open field test |  |  |  |  |
| --- | --- | --- | --- | --- |
| Group | Travel distance (m) | Speed (cm/s) | Center duration in 10 mins (s) | Center duration in the first 5 mins (s) |
| WT+GFP | 4074 ± 800 | 6.79 ± 1.33 | 34.34 ± 19.70 | 12.98 ± 3.51 |
| AD+GFP | 3162 ± 516 | 5.27 ± 0.86 | 81.31 ± 58.17 | 58.55 ± 37.36 |
| AD+4D | 2850 ± 828 | 4.75 ± 1.38 | 42.87 ± 52.83 | 8.76 ± 5.93 |
| Odd ratio of navigational strategy in the learning phase MWM (WT/GFP vs. AD/GFP) |  |  |  |  |
| thigmotaxis | random | chaining | hippocampal |  |
| 0.22 | 0.6 | 0.43 | 3.64 |  |
| **Odd ratio of navigational strategy in the reversal phase MWM (WT/GFP vs. AD/GFP)** |  |  |  |  |
| **thigmotaxis** | **random** | **chaining** | **hippocampal** | **perseverance in reversal phase 1<sup>st</sup> day** |
| 0.3 | 0.36 | 0.3 | 5 | 5.5 |
| **Odd ratio of navigational strategy in the learning phase MWM (AD/4D vs. AD/GFP)** |  |  |  |  |
| **thigmotaxis** | **random** | **chaining** | **hippocampal** |  |
| 0.36 | 0.98 | 1.67 | 1.51 |  |
| **Odd ratio of navigational strategy in the reversal phase MWM (AD/4D vs. AD/GFP)** |  |  |  |  |
| **thigmotaxis** | **random** | **chaining** | **hippocampal** | **perseverance in reversal phase 1<sup>st</sup> day** |
| 0.55 | 1.17 | 0.8 | 1.32 | 4.36 |

To better dissect the nature of these non-significant trends in center-exploratory behavior, we increased the temporal resolution of our analysis by splitting the total duration of the test of 10 minutes into two intervals of 5 minutes each. This revealed that during the initial 5 minutes of the OFT WT/GFP mice spent significantly less time in the center relative to AD/GFP (mean ± SD: 12.9 ± 3.5 s vs. 58.5 ± 37.3 s, *p* < 0.05; Fig. 2C, right). Notably, AD/4D mice significantly reduced the duration of their exploration of the center relative to AD/GFP to values more similar to WT/GFP controls (AD/4D vs. AD/GFP, mean ± SD: 8.7 ± 5.9 s vs. 58.5 ± 37.3 s, *p* < 0.01; AD/4D vs. WT/GFP, *p* = 0.16 Fig. 2C, right).

Interestingly, in addition, the speed of WT/GFP in the center was significantly higher than AD/GFP (mean ± SD: 12.8 ± 2.6 cm/s vs. 6.8 ± 1.0 cm/s, *p* < 0.01, Fig. S2A). AD/4D also increased their speed in center relative to AD/GFP and to levels similar to WT/GFP mice (AD/4D vs. AD/GFP, mean ± SD: 12.9 ± 5.7 cm/s vs. 6.8 ± 1.0 cm/s, *p* < 0.05; AD/4D vs. WT/GFP, *p* = 0.95, Fig. S2A). These differences were no longer detectable among the three groups during the last 5 minutes of the OFT (Fig. S2B) explaining why groups appeared not-significantly different when considering the total time of 10 minutes of the test. Collectively, these observations highlight the importance of dissecting the temporal dynamics of behavior showing that increased neurogenesis by expansion of endogenous NSC partially rescued baseline exploration, particularly during initial phase of the test.

Next, we investigated whether 4D-enhanced neurogenesis was sufficient to rescue spatial learning and memory deficits of AD mice during a navigational cognitive task. While analyzing standard parameters of the MWM tests, i.e.: pathlength and speed, we found that WT/GFP travelled less and were slower in reaching the goal relative to both AD/GFP and AD/4D (pathlength mean = 888, 1,554 and 1,506 cm; speed mean = 17.1, 20.4, 20.6 cm/s, for the three groups, respectively). No difference in neither pathlength nor speed was found when comparing AD/GFP and AD/4D mice.

Despite the lack of differences in pathlength and speed between controls and AD/4D mice, it has been consistently shown that assessment of navigational strategies is essential for to discriminate between egocentric and hippocampal-specific, allocentric navigation and perseverance after reversal (Garthe and Kempermann, 2013; Grech, Nakamura and Hill, 2018) as opposed to pathlength and/or speed. To perform these analyses, navigational strategies were classified based on a previously adopted algorithm used in several previous reports (Berdugo-Vega *et al*., 2020; Overall *et al*., 2020; Schwab *et al*., 2022; Lehr *et al*., 2023) to classify i) thigmotaxis ii) random strategies iii) egocentric chaining or vi) hippocampal-dependent, allocentric directed strategies and, finally, v) perseverance upon reversal (see Material and Methods and Figs. 2D and S2E).

When comparing the navigational strategies during 4 days of learning, WT/GFP mice progressed from thigmotaxis to random strategies and chaining to finally utilize mainly allocentric strategies more efficiently than AD/GFP mice (change in probability of WT/GFP vs. AD/GFP: contextual = 124%, thigmotaxis = -74%; OR WT/GFP vs. AD/GFP: contextual = 3.64, thigmotaxis = 0.21, Wald-test *p* < 0.0001; Fig. 2E, left and S2E). This was expected and corroborates the known impairment in learning characterizing the AD phenotype (Chen *et al*., 2000; Reiserer *et al*., 2007; Baeta-Corral and Giménez-Llort, 2015; Berkowitz *et al*., 2018). Notably, however, AD/4D mice showed an attenuation of the AD phenotypes with a reduced thigmotaxis similar to WT/GFP mice (change in probability of AD/4D vs. AD/GFP: thigmotaxis = -58%; OR AD/4D vs. AD/GFP: thigmotaxis = 0.36, Wald-test *p* < 0.0001) while displaying a trend, although not significant, toward the use of hippocampal-dependent allocentric strategies (change in probability of AD/4D vs. AD/GFP: contextual = 34%; OR AD/4D vs. AD/GFP: contextual = 1.51, Wald-test *p* = 0.06; Fig. 2E, right). After learning, the position of the platform was reversed and re-learning evaluated showing a reduced thigmotaxis and increased perseverance of AD/4D mice (change in probability of AD/4D vs. AD/GFP: thigmotaxis = -38%, perseverance = 187%; OR AD/4D vs. AD/GFP: thigmotaxis = 0.55, Wald-test *p* < 0.05; perseverance = 4.36, Wald-test *p* < 0.01; Fig. 2F, right, S2E-G).

In essence, increasing neurogenesis triggered improvements in some, but not all, aspects of exploratory and navigational behavior and partially restored certain aspects of hippocampal function in the AD brain.

## DISCUSSION

Our study shows that the AD-compromised neurogenic niche is still a susceptible target for genetic expansion of NSC and enhanced AHN despite ongoing neurodegeneration and amyloid plaque load. Most importantly, enhanced AHN alone was sufficient to ameliorate cognitive deficits associated with AD, namely exploratory and navigational behavior. Our findings support the notion that AHN represents a valuable therapeutic target for addressing cognitive impairments in AD.

Adult-born neurons are thought to contribute to hippocampal function by enhancing synaptic plasticity and refining memory precision (Anacker and Hen, 2017; Toda *et al*., 2019; Kempermann, 2022). The observed behavioral recovery in AD/4D mice is consistent with these roles. Importantly, 4D not only drives proliferation of active NSC but also maintains quiescent NSC that are essential for the long-term regenerative potential of the neurogenic niche (Berdugo-Vega *et al*., 2020). This is an important consideration, as sustained neurogenesis is required to support cognitive function throughout life as well as over the course of long-lasting neurodegenerative diseases.

The fact remains that the degree of recovery observed in our study was limited. Spatial learning and memory performance did not fully return to wild-type levels and some behavioral measures remained impaired. In addition, many other aspects of hippocampal function were not investigated, such as long-term memory and mood regulation. Such limited recovery likely reflects the presence of AD-related impairments, including Aβ and tau accumulation, synaptic degeneration, and neuroinflammation (Salta *et al*., 2023) across the whole brain, despite a partially restored hippocampal function. Thus, neurogenesis-based strategies alone are likely insufficient to achieve complete recovery of AD but could still be critical and act synergistically with other therapeutic approaches.

Importantly, cognitive improvements occurred despite unchanged amyloid burden, supporting the emerging view that neurogenic enhancement operates independently from plaque clearance (Marlatt *et al*., 2013). This dissociation has critical therapeutic implications: AHN-targeted interventions could complement amyloid-or tau-directed therapies, potentially providing synergistic benefits through distinct mechanistic pathways. The partial nature of behavioral rescue argues for such combination of approaches rather than neurogenesis-targeted monotherapy.

In summary, expansion of endogenous NSC restores AHN and improves cognitive function in AD mice. Our study highlights the responsiveness of the neurogenic niche despite pathological burden making it a viable target for multi-modal therapeutic strategies to counteract neurodegeneration in AD.

## MATERIALS AND METHODS

### Animal handling

Mice were kept in standard cages with a 12-hour light cycle with water and food ad libitum. Female, 6-month-old, 3xTg AD mice with a mixed C57BL/6, 129X1/SvJ, 129S1/Sv genetic background were used in all experiments. Suspensions of lentiviruses were prepared as previously described at a titer of 10^8-9^ IU/ml (Berdugo-Vega *et al*., 2020) and 1 μl bilaterally injected in the DG of isoflurane-anesthetized mice at a constant flow of 200 nl/minute using a nanoliter-2000 injector (World Precision Instruments) and a stereotaxic frame Model 900 (Kopf Instruments) at ±1.6 mm mediolateral, –1.9 anterior-posterior, and –1.9 mm dorsoventral from bregma. AHN was assessed 12 weeks after surgery by intraperitoneal injection of BrdU (Sigma) dissolved in PBS (50 mg/kg body weight), twice a day for 5 consecutive days. Animals were subsequently subjected to behavioral tests, anesthetized with xylazine/ketamine, and perfused transcardially with saline followed by 4% paraformaldehyde (PFA) fixation in phosphate buffer for immunohistochemistry. Animal procedures were performed according to local regulations (DD 24-9168.11-1/2011-11, TVV 45/2021).

### Behavioral tests

Open field test (OFT) was performed in 60 × 60 cm field with 40 cm high white plywood walls. The room was illuminated with a 100 watts light bulb installed 1.5 m above the field. Each mouse was placed in one corner of the empty arena and allowed to freely explore for 10 minutes. The exploratory activity of the mice was recorded with a camera (Logitech) and analyzed with EthoVision software (Noldus) to assess distance, speed and time spent in the central area (inner 50% open field area). After each trial, the apparatus was cleaned with 75% ethanol to avoid odor cues. Morris Water Maze (MWM) was performed as previously described (Berdugo-Vega *et al*., 2020) in a circular pool (1.89 m diameter) filled with water kept at 19-20°C and made opaque with a white non-toxic pigment. Mice were trained to find the square escape platform (9 × 9 cm) submerged 0.5 cm below the water surface over 8 days including 4-day learning and 4-day reversal. Each day, mice performed 4 trials with a minimum intertrial time of 15 minutes. Each trial lasted a maximum of 2 minutes at the end of which mice were guided to the platform and then placed in individual drying cages equipped with a warm light. On the first day of reversal, the platform was moved to the opposite quadrant to assess re-learning. Data acquisition and analysis were performed with the EthoVision system (Noldus). Navigational strategies were assessed by unbiased, automatic software analysis (Rtrack) of the swimming trajectories according to Overall et al., 2020 with each trial being assigned one predominant strategy among thigmotaxis, random (circling, random search, scanning), chaining, hippocampal (directed, focal, direct) or perseverance, as previously reported (Garthe, Behr and Kempermann, 2009; Garthe and Kempermann, 2013).

### Immunohistochemistry and cell quantification

Brains were harvested after perfusion and post-fixed in 4% PFA at 4°C overnight. After washing with PBS and embedding in 3% low-melting agarose, brains were coronally cut into 40 μm thick slices by a vibratome. Serial sections along the rostral-caudal axis of the hippocampus (every sixth section in one tube, total 6 tubes) were collected and stored in cryoprotectant solution (25% Ethanol and 25% glycerol in 1x PBS) at –20°C. For immunolabeling, slices were washed with PBS, blocked and permeabilized with 10% donkey serum in 0.3% Triton X-100 in PBS for 1.5 hours at room temperature. Primary and secondary antibodies (Supplementary Table 1) were diluted in 3% donkey serum in 0.3% Triton X-100 in PBS and incubated overnight at 4°C or 2 hours at room temperature, respectively, with DAPI being added for the last 10 minutes. For BrdU or Aβ detection, slices were exposed to HCl 2M for 25 minutes at 37°C or 88% formic acid for 3 minutes at 70°C before blocking, respectively. Pictures were acquired with Zeiss ApoTome with maximal intensity projections of three optical sections (10 μm total) or AxioScan microscopes (Carl Zeiss) followed by Extended Depth of Field Calculation. Cell counts were obtained using Photoshop CS5 (Adobe) and analysis of Aβ load performed with Fiji (ImageJ).

### Statistical analysis

Cell count and Aβ quantification were performed on at least 3 mice as biological replicates with data reported as mean ± SD (Table 1 and Supplementary Table 2 and significance calculated by two-tailed unpaired Student’s t-test. A minimum of 5 mice were used for behavioral analyses as indicated in the respective figure legends and data depicted as whisker box (showing median, 25th-75th percentiles and max-min range for travel distance, duration, speed, and perseverance), box plots (proportional change in strategies between groups, and number of mouse performing perseverance), or area charts (navigational strategies). Values of the behavioral tests were reported in Table 1, Table 2, Supplementary Table 2 and Supplementary Table 3. Significance was calculated by i) 2-way ANOVA (evolution of performance over consecutive trials/days), ii) two-tailed unpaired Student’s t-test (difference of performance in the behavioral analysis), or iii) Wald-test of odd ratios assessed by logistic regression (navigational strategies). Analyses were performed using GraphPad Prism or Microsoft Excel.

## CONFLICT OF INTEREST

The authors declare that the research was conducted in the absence of any commercial or financial relationships that could be construed as a potential conflict of interest.

## FUNDING

This work was supported by EU-H2020 Marie Skłodowska-Curie grant (813851) and the Medical Faculty of TU-Dresden and the CRTD.

## ACKNOWLEDGMENTS

The authors thank the CRTD animal facility for their assistance, Dr. Gabriel Berdugo-Vega for technical support, and members of the lab for feedback throughout the project. The authors declared and acknowledged that Generative AI tools (ChatGPT and Claude) were used to assist of refinement of language usage in the manuscript.

## SUPPLEMENTARY FIGURES

**Supplementary Figure 1.**
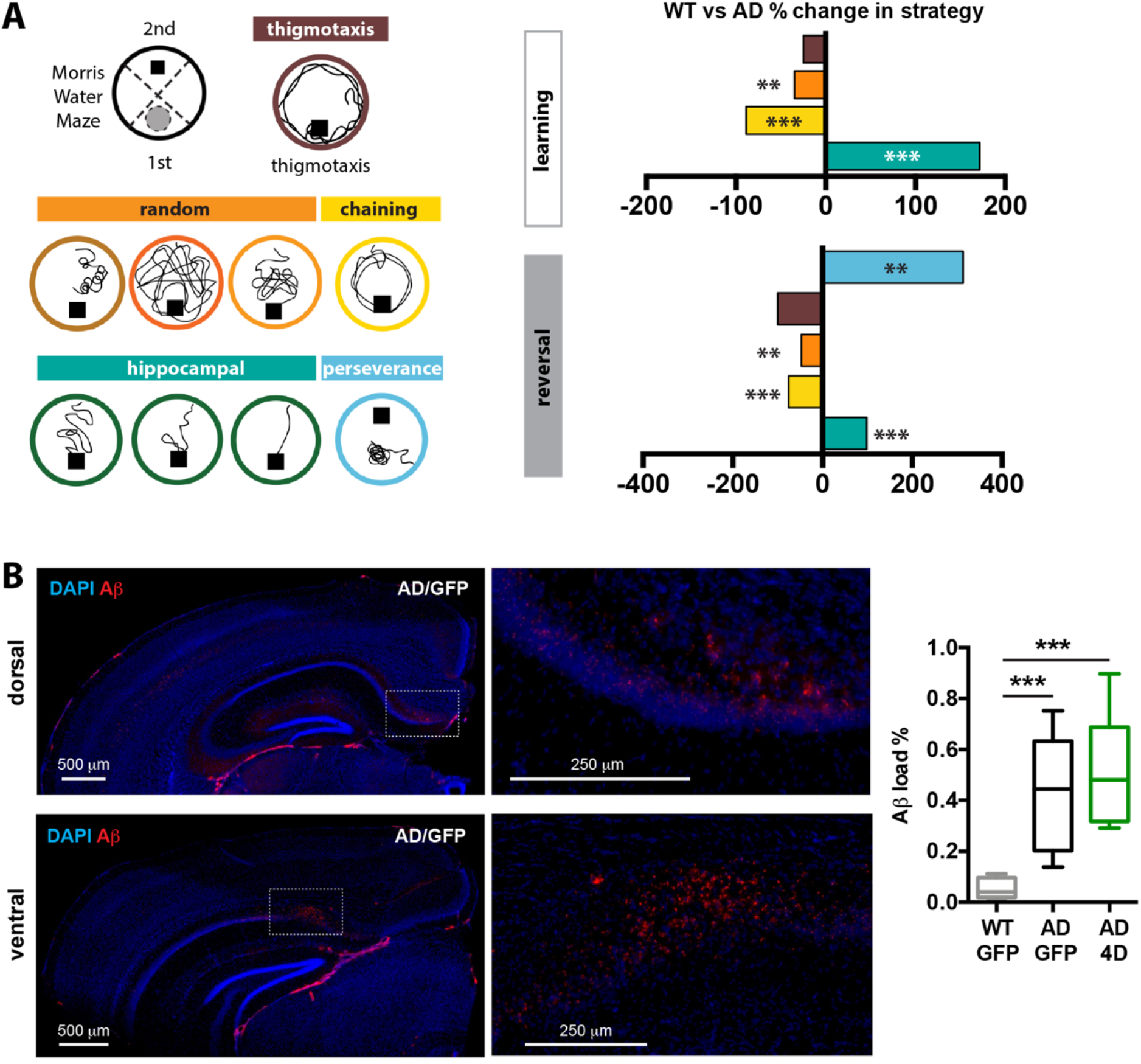
Analysis of AD phenotypes and pathologies in 3xTg AD mice. **A**) Searching strategies assigned to each trial by the algorithm in MWM. Comparison of the percentage change in searching strategies across trials during the learning and reversal phase between WT and AD at 6-month-old. (n = 8 and 6 for WT and AD groups respectively), Wald-test, ** *p* < 0.01, *** *p* < 0.001. **B**) Aβ staining picture and quantification of Aβ levels in the hippocampus. Values represent whisker box (n = 5, 10, and 10 for WT, AD/GFP, and AD/4D groups respectively), unpaired Student’s t-test, *** *p* < 0.001. Values of mean ± SD are listed in Supplementary Table 2.

**Supplementary Figure 2.**
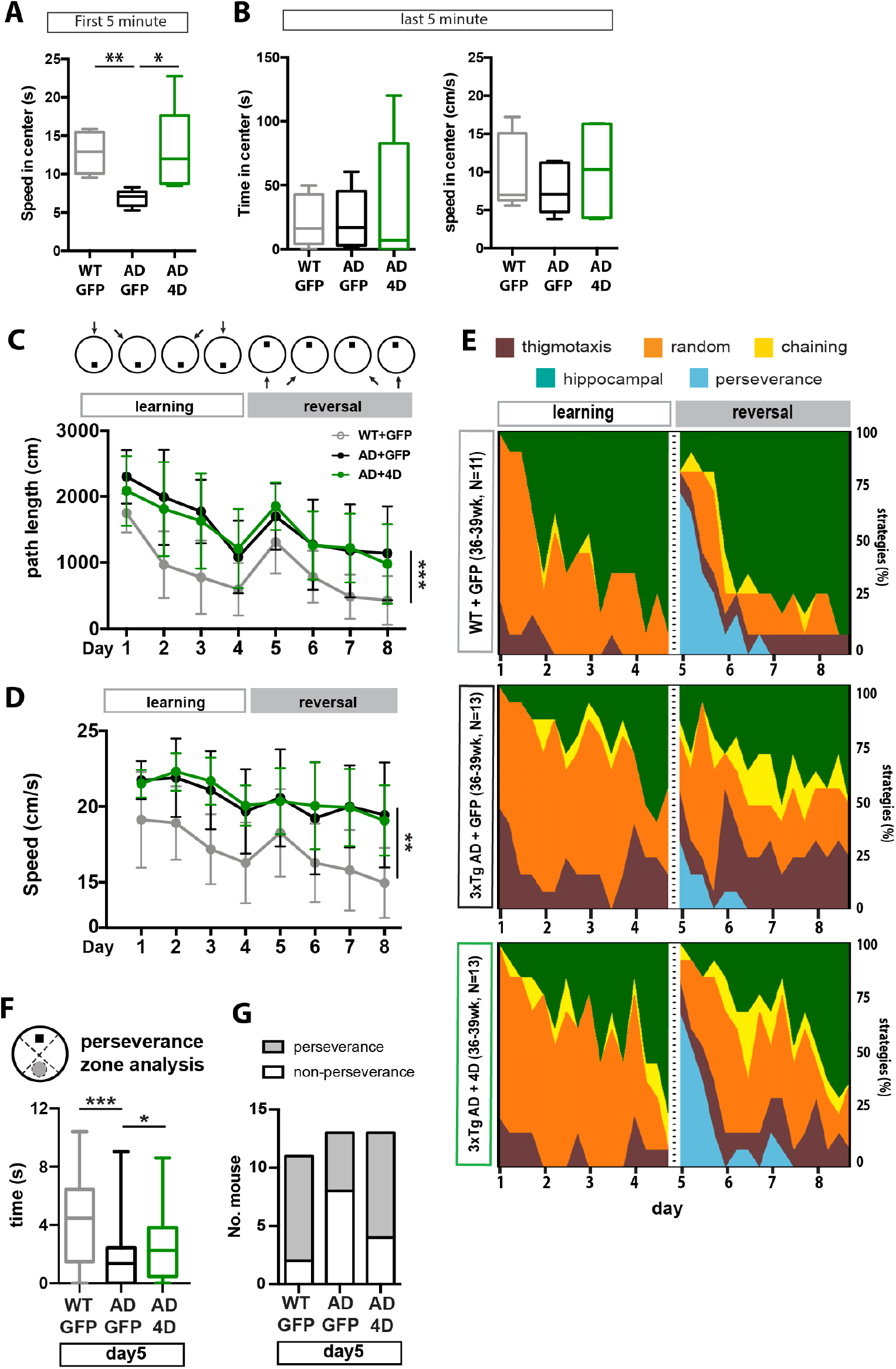
Additional analysis in navigational performance of WT, AD/GFP and AD/4D mice. **A**) Speed in the center (cm/s) of the open field during the initial 5-minute exploration. **B**) Time (s) and speed (cm/s) in the center of the open field during the last 5-minute exploration. **C**) Depiction of starting points in MWM and swimming path length (cm) in MWM. A significant difference was observed between WT and AD/GFP (two-way ANOVA, time F (7, 154) = 25.39, *p* < 0.0001, group F (1,22) = 20.74, *p* < 0.001, interaction F (7, 154) = 1.890, *p* = 0.07). No difference was between D/4D and AD/GFP (two-way ANOVA, time F (7, 168) = 22.09, *p* < 0.0001, group F (1, 24) = 0.08419, *p* = 0.77, interaction F (7, 168) = 0.6700, *p* = 0.69). **D**) swimming speed (cm/s) in MWM. A significant difference was observed between WT and AD+GFP (two-way ANOVA, time F (7, 154) = 11.31, *p* < 0.0001, group F (1, 22) = 13.91, *p* < 0.001, interaction F (7, 154) = 1.109, *p* = 0.36). No difference was between D/4D and AD/GFP (two-way ANOVA, time F (7, 168) = 8.053, *p* < 0.0001, group F (1, 24) = 0.05675, *p* = 0.081, interaction F (7, 168) = 0.3622, *p* = 0.92). **E**) Schematics, color code, and the contribution of swimming strategies as percentages. **F**) perseverance zone analysis on fifth day of MWM. **G**) Number of mice performing perseverance at day 5 within the group. **A**) – **B**) Values represent whisker box (n = 6, 5, and 6 for WT, AD/GFP, and AD/4D groups respectively), unpaired Student’s t-test, * *p* < 0.05, ** *p* < 0.01. **C**) – **G**) For WT, AD/GFP, and AD/4D groups, the numbers of mice were indicated (n=11, 13, and 13 respectively). Values represent whisker box. Unpaired t test, * *p* < 0.05; ** *p* < 0.01; *** *p* < 0.001. Values are listed in Supplementary Table 3.

## SUPPLEMENTARY TABLES

**Supplementary Table 1.** Primary and secondary antibodies for immunohistochemistry.

| Primary antibodies |  |  |  |
| --- | --- | --- | --- |
| Antigen | Dilution | Supplier | Catalog number |
| BrdU | 1:250 | Abcam | ab6326 |
| Sox2 | 1:500 | Santa Cruz Biotechnology | Sc-17320 |
| S100 $\beta$ | 1:1000 | Abcam | ab14688 |
| DCX | 1:100 | Abcam | ab18723 |
| NeuN (Fox3) | 1:1000 | Abcam | ab104225 |
| GFP | 1:1000 | Abcam | Ab13970 |
| Aβ | 1:1500 | Novus Biologicals | NBP2-13075 |
| **Secondary antibodies:** IgG raised in donkey (against goat, mouse, rabbit, rat and guinea pig), conjugated to different fluorophores (DyLight, Alexa or Cyanines), all purchased from Jackson Immunoresearch. |  |  |  |

**Supplementary Table 2.** Comparison of navigational performance and amyloid plaque load in the hippocampus.

| Odd ratio of navigational strategy in the learning phase MWM (WT vs. AD) |  |  |  |  |
| --- | --- | --- | --- | --- |
| Thigmotaxis |  | Random | Chaining | Hippocampal |
| 0.75 |  | 0.46 | 0.09 | 5.22 |
| Odd ratio of navigational strategy in the reversal phase MWM (WT vs. AD) |  |  |  |  |
| Thigmotaxis | Random | Chaining | Hippocampal | Perseverance in reversal phase 1<sup>st</sup> day |
| 0 | 0.43 | 0.18 | 5.21 | 5.6 |
| Hippocampal Aβ load (%) |  |  |  |  |
| WT+GFP | 0.05195 ± 0.04213 |  |  |  |
| AD+GFP | 0.4308 ± 0.2218 |  |  |  |
| AD+4D | 0.5094 ± 0.1970 |  |  |  |

Odd ratios (OR) of strategy comparing WT and 3xTg AD mice and hippocampal Aβ load (%), corresponding to Fig S1.

**Supplementary Table 3.**
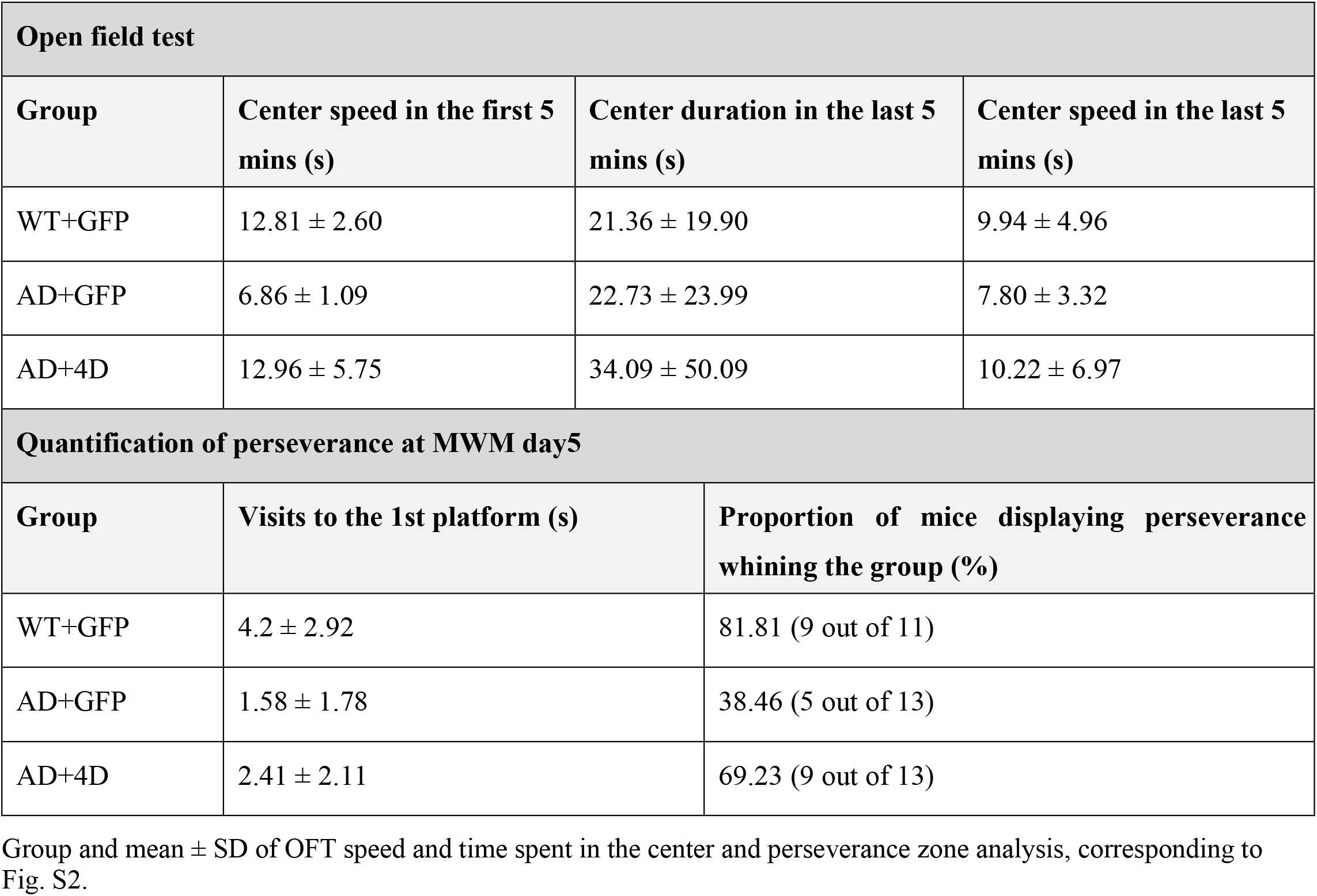
Comparison of navigational performance and amyloid plaque load in the hippocampus.

## Notes

### Competing Interest Statement

The authors have declared no competing interest.

